# Establishing a causal role for left ventrolateral prefrontal cortex in value-directed memory encoding with high-definition transcranial direct current stimulation

**DOI:** 10.1101/2023.01.04.522746

**Authors:** Linfeng Tony Han, Michael S. Cohen, Liqin Ken He, Laura M. Green, Barbara J. Knowlton, Alan D. Castel, Jesse Rissman

**Affiliations:** Department of Psychology, University of California, Los Angeles, Los Angeles, CA 90095; Department of Psychology, University of Pennsylvania, Philadelphia, PA 19104; Department of Psychology, Tsinghua University, Beijing, China 100084; Department of Psychology, University of Chicago, Chicago, IL 60637; Department of Psychiatry and Biobehavioral Sciences, University of California, Los Angeles, Los Angeles, CA 90095

**Keywords:** Verbal memory, value-directed remembering, selective encoding, HD-tDCS, ventrolateral prefrontal cortex, hemispheric lateralization

## Abstract

One critical approach for promoting the efficiency of memory is to adopt selective encoding strategies to prioritize more valuable information. Past neuroimaging studies have shown that value-directed modulation of verbal memory depends heavily on engagement of left-lateralized semantic processing regions, particularly in ventrolateral prefrontal cortex (VLPFC). In the present study, we used high-definition direct current stimulation (HD-tDCS) to seek evidence for a causal role of left VLPFC in supporting the memory advantage for high-value items. Three groups of healthy young adult participants were presented with lists of words to remember, with each word accompanied by an arbitrarily assigned point value. During the first session, all participants received sham stimulation as they encoded five lists of 30 words each. Two of these lists were immediately tested with free recall, with feedback given to allow participants to develop metacognitive insight and strategies to maximize their point total. The second session had the exact same structure as the first, but the groups differed in whether they received continued sham stimulation (*N*=22) or anodal stimulation of the left VLPFC (*N*=21) or right VLPFC (*N*=20). Those lists not tested with immediate recall were tested with recognition judgments after a one-day delay. Since no brain stimulation was applied during this Day 2 test, any performance differences can be attributed to the effects of stimulation on Day 1 encoding processes. Anodal stimulation of left VLPFC significantly boosted participants’ memory encoding selectivity. In comparison, no such effect was seen in participants who received right VLPFC or sham stimulation. Estimates of recollection- and familiarity-based responding revealed that left VLPFC stimulation specifically amplified the effects of item value on recollection. These results demonstrate a causal role for left VLPFC in the implementation of selective value-directed encoding strategies, putatively by boosting deep semantic processing of high-value words. Our findings also provide further evidence on the hemispheric lateralization of value-directed verbal memory encoding.

## 1. Introduction

Humans tend to prioritize the learning of information that is relatively important at the expense of less important information, especially when faced with a large amount to learn in a limited time. For example, while people may find it impossible to memorize where all their personal items are located, they usually tend to prioritize remembering the locations of items deemed to be more valuable (e.g., laptop, phone, keys). The ability to remember things selectively based on value allows one to maximize the efficiency and utility of learning (Castel et al., 2002, 2007). This process may rely on goal-directed memory strategies as well as on more automatic memory processes (Castel, 2022).

Early insights into the neural substrates of value-based memory in humans came from fMRI work demonstrating that increased activity in the dopaminergic midbrain, including the ventral tegmental area (VTA) and nucleus accumbens, is associated with better memory encoding of high-value items (Adcock et al., 2006). They also found these critical regions of the brain’s so-called ‘reward system’ increased their functional connectivity with the hippocampus during exposure to valuable items, which predicted enhanced subsequent memory for these items. Although fMRI data cannot directly index neurotransmitter levels nor causal interactions, these findings suggest a reward-based learning pathway where dopamine release in the hippocampus potentiates the learning of high-value items. However, memory selectivity for high-value items putatively driven by this pathway does not always emerge on immediate testing but rather is most apparent after a delay, typically of 24-hours or longer (Murayama & Kuhbandner, 2011; Spaniol et al., 2014). This implies that the dopaminergic midbrain reward system especially impacts the efficacy of hippocampal-dependent memory consolidation processes that take hours to sculpt synaptic connections following initial encoding.

Compared with dopaminergic reward-based learning, another critical mechanism for the encoding of high-value items is the engagement of metacognitive control and selective memory encoding strategies. In a value-directed remembering (VDR) task (Castel et al., 2002; Castel, 2008; see Knowlton & Castel, 2022, for a recent review), participants study multiple lists of words assigned with different point values. As they are tested and given immediate feedback after each list, they become increasingly selective in later word lists. Such a rapid change in encoding selectivity is not likely to be a consequence of dopamine-driven reward-based learning (which seems to impact memory over a longer time scale). Instead, it may be driven by more effective learning and subsequent recollection of high-value items due to better encoding strategies (Hennessee et al., 2017, 2019). With task experience, participants learn to modify their encoding strategies to direct more cognitive resources to high-value words across the study-test cycles (Cohen et al., 2017; Murphy et al., 2021). Several fMRI studies have used variants of the VDR paradigm and revealed that activation in the left ventrolateral prefrontal cortex (VLPFC) and left posterior lateral temporal is associated with memory selectivity, which demonstrates a lateralized neural mechanism for value-based memory enhancement that is separate from the midbrain reward system (Cohen et al., 2014, 2016). Importantly, these left-lateralized cortical regions are known to be heavily involved in semantic processing (Jackson, 2021; Vigneau et al., 2006), which is consistent with participants’ use of verbal elaboration strategies to facilitate the learning of high-value words (e.g., reflecting deeply on the meaning or self-relevance of the to-be-remembered words, or even linking the words together to make up a memorable story). Though these fMRI findings certainly suggest a relationship between strategic engagement of deep semantic processing and the selective encoding of high-value items, it remains unclear whether putative semantic processing regions like left VLPFC are causally involved in the value-incentivized memory encoding process.

In the present study, we used transcranial direct current stimulation (tDCS) to test whether experimentally augmented cortical activity in the left VLPFC would result in higher memory encoding selectivity. tDCS is a neuromodulation technique that regulates cortical excitability in a targeted brain region, and it has been demonstrated to be effective for intervening with psychiatric disorders in clinical populations (Kekic et al., 2016) and enhancing various cognitive functions in healthy individuals, including those functions specific to memory (Galli et al., 2019; Hill et al., 2016; Mancuso et al., 2016). tDCS has also been considered as a useful technique for establishing the causal involvement of a given brain region in a specific behavior of interest (Reinhart et al., 2017). The effects of tDCS are generally thought to be influenced by stimulation polarity, such that regions under the anode are depolarized and thus facilitated whereas regions under the cathode are hyperpolarized and thus inhibited (Nitsche & Paulus, 2000); however, the inhibitory consequences of cathodal stimulation in cognitive tasks is more controversial and inconsistently observed (Jacobson et al., 2012). Whereas conventional tDCS delivers electrical current that flows between two large pad electrodes (∼30 cm^2^) resulting in a broad profile of stimulation, high-definition tDCS (HD-tDCS) is a more focal technique that allows for targeted stimulation of a smaller brain region (Alam et al., 2016; Datta et al., 2009). Thus, we elected to use HD-tDCS to concentrate the current intensity with greater anatomical precision.

We targeted our stimulation site as the left VLPFC given its critical role in semantic processing and encoding of valuable verbal information (Cohen et al., 2014, 2016; Thompson-Schill et al., 1997). In addition, we included two control groups: one group received stimulation targeting the homologous region in the right hemisphere (right VLPFC) and one group received sham stimulation. We hypothesized that active anodal stimulation of the left VLPFC would result in a larger enhancement effect in memory encoding selectivity compared with anodal stimulation of the right VLPFC, which would demonstrate the hemispheric lateralization (Benson & Zaidel, 1985) of this causal effect. The inclusion of a sham group provides an additional baseline, allowing us to test the possibility that any VLPFC stimulation is beneficial for encoding selectivity. However, given the strong left-lateralization apparent in our prior fMRI studies, we predicted that only the left VLPFC stimulation group would show benefits relative to sham. Importantly, we also designed our experiment in such a way so as to allow each participant to serve as their own control. By beginning every participant’s encoding session with sham stimulation and then assigning them (in a double-blind fashion) to one of the three experimental conditions (left VLPFC stimulation, right VLPFC stimulation, or additional sham stimulation) for the second half of the encoding session, we were able to conduct within-subjects contrasts that inherently control for individual differences in memory ability and strategy use, and then we could conduct between-subjects analyses to examine effects of group assignment on these difference scores.

In our experiment, we adapted the VDR paradigm to gauge the consequences of HD-tDCS on verbal memory encoding, with a focus on how stimulation impacts participants’ ability to prioritize the memorization of high-value words. As we have done in prior work, value was operationalized as the number of points that could be earned for successfully remembering each word. The experiment was administered over two consecutive days, with HD-tDCS stimulation always occurring on the first day during encoding. By testing memory retention on the second day, we ensured that stimulation effects were no longer influencing brain activity during retrieval, allowing us to more cleanly isolate the impact on encoding. As tDCS effects are known to linger for minutes to hours after the stimulation ends (Kuo et al., 2013; Nitsche & Paulus, 2001; Stagg et al., 2013), retrieval processes evoked during immediate memory testing would likely be impacted by stimulation. Our protocol did in fact include immediate free recall tests on a subset of the word lists, and participants were instructed to be prepared for the possibility of such testing on all lists. When tested with free recall, participants were given detailed feedback on their performance, as this is known to provide helpful metacognitive insights that encourage more strategic encoding on the ensuing lists (Cohen et al., 2017). But by omitting free recall testing after some of the lists, we were able to save our assessment of value-based memory effects until the following day when we could be sure that retrieval processes would be unaffected by the stimulation. Participants’ subsequent recognition memory for the words on these untested lists was thus the critical measure of interest in our experiment.

Assuming that anodal stimulation of the left VLPFC would enhance memory encoding selectivity, we also examined whether any increase in selectivity would be most apparent in words that were recollected, or whether both recollection and familiarity would be affected. Some prior studies have reported that selective strategic processing benefits both recollection and familiarity on later recognition tests (Cohen et al., 2017), while others have found that value at encoding primarily benefits recollection (Elliott et al., 2020; Hennessee et al., 2017). We adopted the dual-process signal-detection (DPSD) model to dissociate recollection and familiarity based on receiver operating characteristic (ROC) curves derived from participants’ recognition confidence ratings, and examined if HD-tDCS affected one or both of these memory processes.

## 2. Methods

### 2.1 General Design

We administered anodal or sham HD-tDCS stimulation to either left or right VLPFC under a double-blind protocol, which resulted in participants being assigned to one of four stimulation groups: left-anodal, right-anodal, left-sham, and right-sham stimulations. We combined the latter two groups of participants who received sham stimulation into a homogenous group throughout the data analysis procedure as we did not expect sham stimulation of contralateral sites to make any difference. As strategic semantic encoding has been demonstrated to be a highly left-lateralized process (Cohen et al., 2014, 2016), we expected enhanced memory encoding selectivity induced by anodal stimulation of the left VLPFC (Group A, Fig. 1A) but not anodal stimulation of the right VLPFC (Group B, Fig. 1A) or sham stimulation (Group C, Fig. 1A). Anodal stimulation is generally thought to have an enhancing effect on cognition by boosting the excitability of cortical neurons in the targeted region (Nitsche & Paulus, 2000). In HD-tDCS protocols, this is typically achieved by placing a single anodal electrode over the targeted region and surrounding it by a ring of several cathodal electrodes (Alam et al., 2016). Although it is technically possible to reverse the polarity of this montage and surround a cathodal electrode with a ring of several anodal electrodes, we did not include a cathodal stimulation group because prior studies have suggested that cathodal stimulation can either suppress or enhance cortical function in unpredictable and often nonlinear ways (Batsikadze et al., 2013; Brückner & Kammer, 2017; Shilo & Lavidor, 2019). Indeed a meta-analysis of tDCS studies concluded that cathodal stimulation of non-motor regions rarely causes inhibition (Jacobson et al., 2012). Given the uncertain outcomes associated with cathodal stimulation, we focused on whether we could enhance value-related memory selectivity with anodal stimulation, and whether this effect would be greater when stimulation was applied over left VLPFC versus right VLPFC.

**Figure 1.**
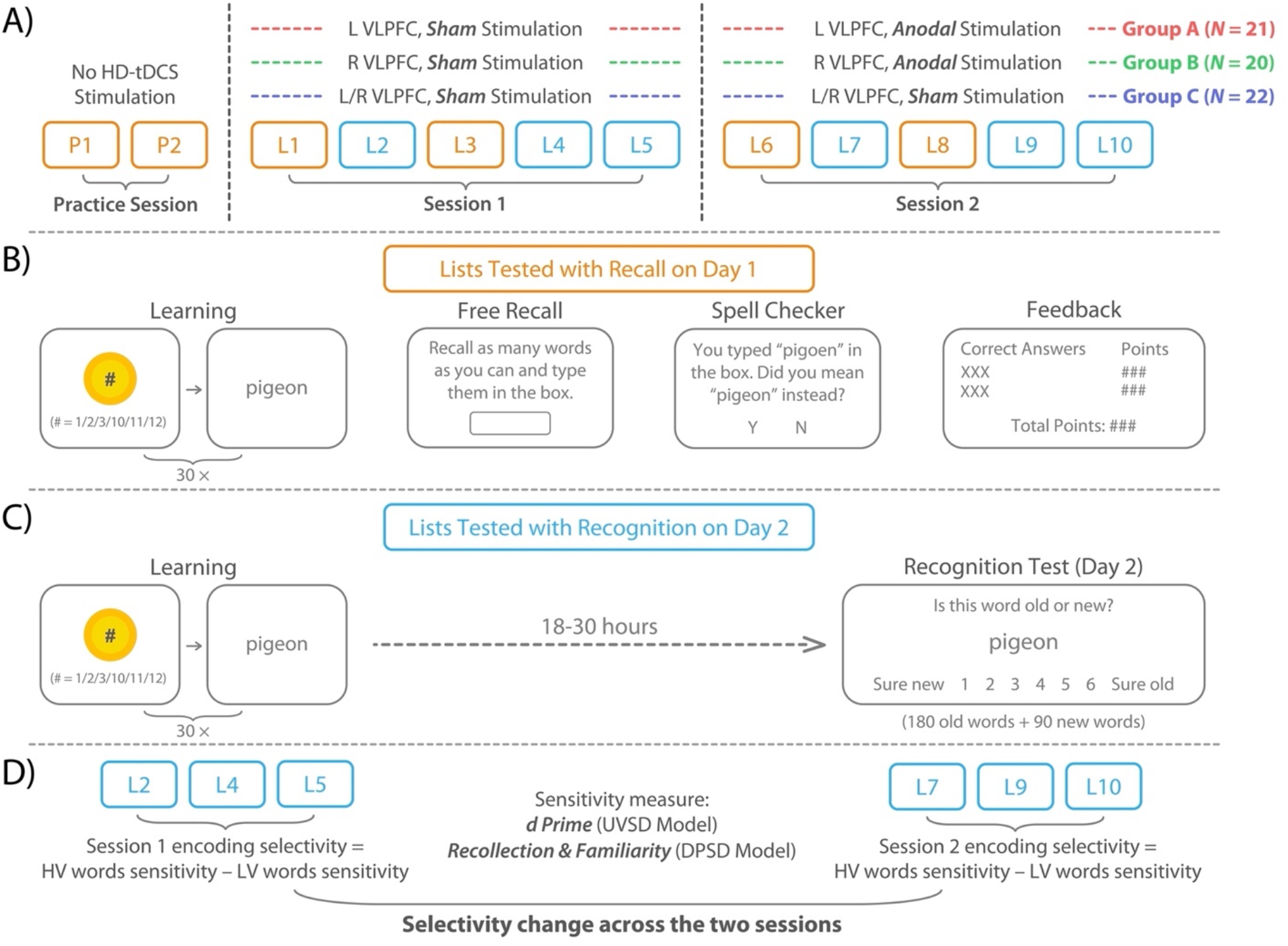
Experimental Procedures and Data Analysis Framework. **A) Overall Day 1 list learning and procedure and stimulation protocol**. All participants completed a practice session (no stimulation) and Session 1 (sham stimulation), after which HD-tDCS stimulation parameters varied by group assignment for Session 2 (Group A: anodal stimulation of left VLPFC; Group B: anodal stimulation of right VLPFC; GROUP C: sham stimulation). Sessions 1 and 2 were separated by only a few minutes and had exactly the same learning and testing structure, with a subset of the lists tested with immediate free recall (orange) and other lists not tested until Day 2 (blue). **B) Learning and testing procedures for the lists tested on Day 1**. Each 30-word list was presented one word at a time, with each word preceded by a point value displayed on a golden coin, indicating how many points (1, 2, 3, 10, 11, or 12) could be earned for later remembering this word. Lists P1, P2, L1, L3, L6, and L8 were tested with immediate free recall on Day 1, and participants received feedback on their performance. **C) Learning and testing procedures for the lists tested on Day 2**. Lists L2, L4, and L5 (from Session 1) and L7, L9, and L10 (from Session 2) were not tested with immediate recall on Day 1 but rather were tested on Day 2 (after 18-30 hours) with judgments of recognition confidence on a 1 to 6 scale. **D) Indexing changes in value-related memory encoding selectivity**. The difference between the recognition memory sensitivity—measured either by *d’* (UVSD model) or recollection and familiarity estimates (DPSD model)—for high-value and low-value words was calculated separately for words that had been learned in each session. This value-related difference score was designated as the encoding selectivity for the session, and the encoding selectivity change across the two sessions was calculated by subtracting the encoding selectivity in Session 1 from that in Session 2.

Importantly, while we manipulated the stimulation type (left-anodal, right-anodal, or sham) as a between-subject factor, we added a within-subject comparison to minimize the impact of individual differences in task performance. Specifically, we had all participants go through a no-stimulation practice session and then a sham-stimulation experimental session (Session 1) prior to undergoing anodal stimulation or continued sham stimulation (Session 2). This protocol allowed participants to develop some encoding strategies and reach relatively high selectivity before they received different types of stimulation, hence enabling us to gauge each individual’s baseline performance and evaluate the effects of different types of HD-tDCS stimulation by measuring the selectivity change across the two testing sessions in a within-subject fashion.

### 2.2 Participants

Our experiment involved three groups of participants (left-anode, right-anode, and sham), with the latter two groups established as control groups. As we hypothesized that the anodal stimulation of the left VLPFC would significantly boost encoding selectivity relative to the other two groups, we conducted an *a priori* power-analysis (Faul et al., 2007) for a one-tailed independent-sample *t*-test. A target sample size of 26 and 21 participants per group was suggested when we assumed 80% power and an effect size of Cohen’s *d* = 0.7 and 0.8, respectively. Based on this, we recruited a total of 72 participants (24 per group) from the University of California, Los Angeles undergraduate student community. Data from 9 participants were excluded for extreme memory performance or lost or damaged data (see the Data Exclusion section). Due to the COVID-19 pandemic that commenced in early 2020 and prevented additional data collection, we were not able replace excluded participants to fulfill our ideal target sample size. Thus, our final dataset consists of 63 participants’ data, with 21 in the left-anode group (*M*_*age*_ = 20.4, *SD* = 1.17), 20 in the right-anode group (*M*_*age*_ = 19.7, *SD* = 1.22), and 22 in the sham group (11 left-sham and 11 right-sham, *M*_*age*_ = 20.6, *SD* = 3.27). We considered this sample size acceptable given that similar or smaller sample sizes have been used in prior studies investigating tDCS modulation of memory (Huang et al., 2021).

### 2.3 High-Definition Transcranial Direct Current Stimulation

We administered high-definition transcranial direct current stimulation (HD-tDCS) via the Soterix (Soterix Medical Inc., New York, NY, USA) 1×1 low-intensity stimulator (Model 1300A) coupled with the 4×1 multichannel stimulation interface (Model 4×1-C3A). The tDCS was applied through a set of five ring-shaped HD electrodes (each 1.2 cm diameter), which were snapped into plastic electrode holders that were fitted into a fabric head cap with holes positioned according to the standard 10-10 EEG layout. The electrode holders were filled with electro-conductive gel (Soterix HD-GEL). For each active stimulation session, the current intensity was quickly raised from 0 to 2 milliamperes (mA), and a constant current of 2 mA was applied at the anode for a 20 min duration; four cathodes served as return electrodes with the load evenly divided between them. For each sham stimulation session, the current intensity was briefly ramped up to 2 mA at the onset of the session (to create the physical sensation of stimulation) and then promptly ramped down to a minimum level (approximately 0.01 mA); this ramp up/down process was repeated at the tail end of the 20 min session. We selected a stimulation intensity of 2 mA because this dosage is commonly used in HD-tDCS studies, including memory studies involving prefrontal stimulation (e.g., Nikolin et al., 2015) and is known to be well-tolerated (Reckow et al., 2018).

For stimulation of the left VLPFC, we placed the anode at F5 and the four return cathodes at C1, C5, F9, and Fp1, respectively (Figs. 2A and 2B for 2D and 3D views of the placement of electrodes). The selection of this montage was based on modeling conducted using Soterix Medical’s HD-Explore™ neurotargeting software (http://soterixmedical.com/software/hd-explore) with the goal of maximizing current flow over the specific region of left VLPFC (Figs. 2C and 2D for 3D and 2D illustration of the simulated field intensity) that both exhibited peak value-related fMRI activation effects during word encoding in a value-directed-remembering task (Cohen et al., 2014, 2016) and showed strong activation associated with semantic processing in an automated meta-analysis of fMRI studies (https://neurosynth.org). For stimulation of the right VLPFC, we created a precisely homologous montage by placing the anode at F6 and the four cathodes at C2, C6, F10, and Fp2.

**Figure 2.**
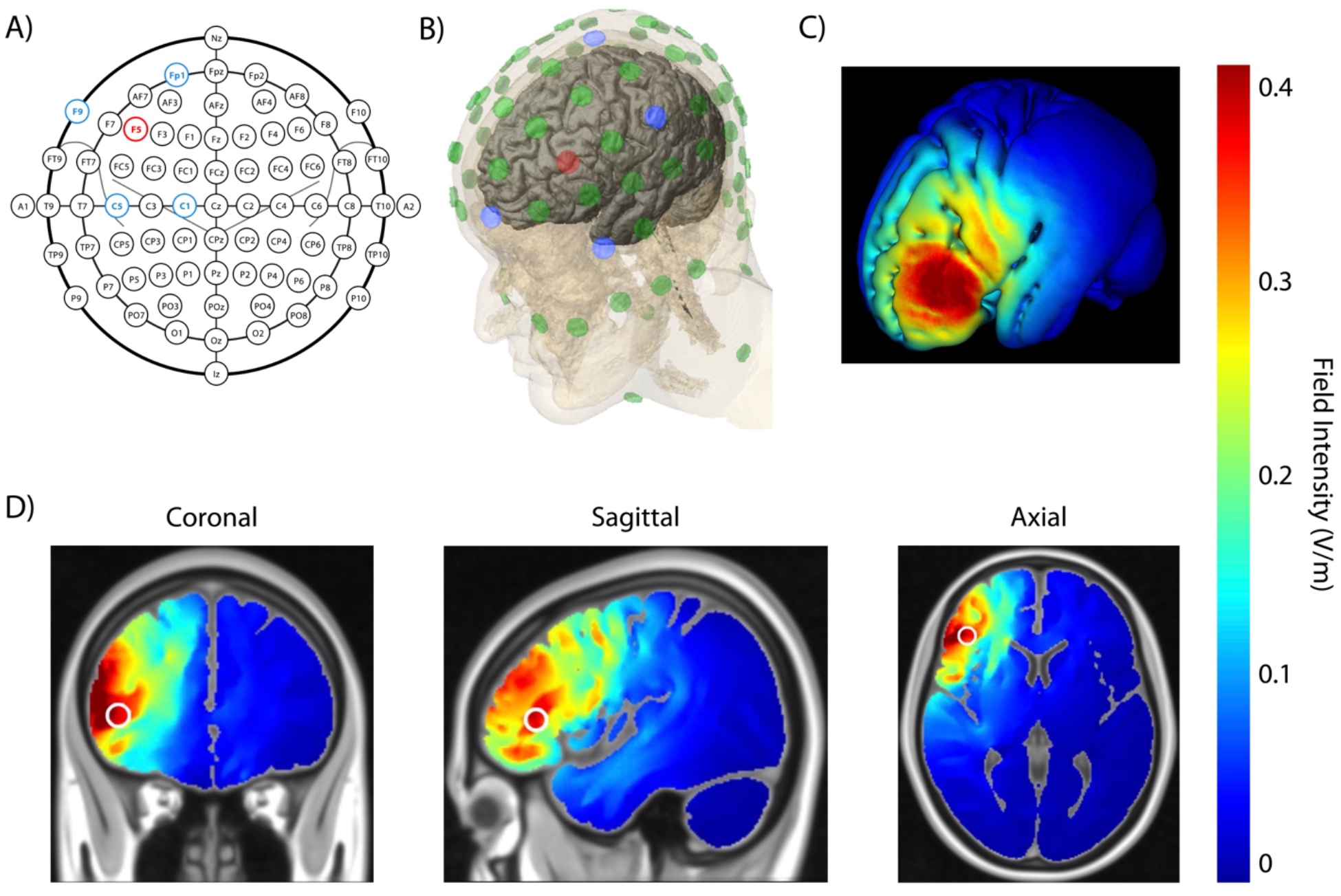
High-Definition Transcranial Direct Current Stimulation. **A) 2D illustration of the electrode placement on a standard EEG cap**. For stimulation of the left VLPFC, we placed the anode at F5 (red) and the four return cathodes at C1, C5, F9, and Fp1 (blue). For stimulation of the right VLPFC, we created a precisely homologous montage by placing the anode at F6 and the four cathodes at C2, C6, F10, and Fp2. **B) 3D illustration of the electrode placement**. The placement of the anode (red) and cathodes (blue) for stimulation of the left VLPFC is shown over a template brain. **C) 3D illustration of computationally modeled current flow intensity**. The simulated field intensity (as rendered by HD-Explore) is shown for 2 mA stimulation targeting left VLPFC. **D) 2D illustration of computationally modeled current flow intensity**. Coronal, sagittal, and axial views of the simulated field intensity in the brain for 2 mA stimulation targeting left VLPFC.

### 2.4 Materials and Procedure

We adopted a similar word selection protocol used in prior behavioral and fMRI studies of value-directed remembering (Cohen et al., 2014, 2016, 2017). We selected 450 four to eight-letter nouns from Clusters 5-8 of the Toglia and Battig (1978) word norms (also see Toglia, 2009) under the criteria that all words must have high familiarity ratings (5.5-7 on a 1-7 scale), moderate to high concreteness ratings (4-6.5), moderate to high imagery ratings (4-6.5), and moderate pleasantness ratings (2.5-5.5).

The experiment was carried out over two consecutive days in a double-blind fashion. This was achieved by ensuring that the experimenter who corresponded to the participant was kept blind to the stimulation type (anodal or sham) because a different experimenter was in charge of operating the HD-tDCS device, and the device was covered to prevent the blind experimenter and participant from viewing its settings.

On Day 1, participants were instructed that they would study and get tested on multiple lists of words while undergoing mild electrical current stimulation of their scalp. They studied twelve lists in total, with each list consisting of 30 words. During the learning phase, each word was presented for 5 s on the center of the screen, preceded by an arbitrarily assigned point value presented for 1.5 s and then a 500-ms blank screen. In each list, half of the words were assigned low values (1, 2, and 3) and the other half were assigned high values (10, 11, and 12).

The first two lists were practice lists (referred to as P1 and P2) that helped the participants understand the learning and testing procedure and provided them with an opportunity to develop some encoding strategies prior to beginning the main experiment. Participants did not undergo any HD-tDCS stimulation during the practice session. Each practice list was immediately followed by a 1.5-min free recall test, where participants were instructed to type as many of the learned words as they could remember into the box presented on the screen, with the goal to maximize their total points. After the free recall test ended, participants went through an automated spellcheck procedure. For each word that the computer flagged as being misspelled (if any), participants could either accept a suggested correction or retype the word. Finally, participants were given feedback on their correctly recalled words, the points associated with each of these words, the total points they obtained on that list, and the highest point total they got so far. The correctly recalled words and their associated points were visually organized such that the low-value words were always presented on the left side of the screen and the high-value words on the right side. In this way, participants were able to see the relative ratio of the correctly recalled low-value versus high-value words, and could then evaluate the effectiveness of their encoding strategies and modify them accordingly to maximize the points earned on later tests.

The ten lists following the two practice lists (referred to as L1-L10) were grouped into two experimental sessions: L1-L5 (referred to as Session 1) and L6-L10 (Session 2). In each session, participants learned the lists while undergoing online stimulation (i.e., concurrent with task performance). We chose online relative to offline stimulation to ensure that our learning phase had a reasonable duration and that the interval between the two sessions was relatively short. The two sessions were separated only by a 2-min break during which the HD-tDCS stimulation parameters could be adjusted. For all participants, we set up the HD-tDCS device before they began Session 1 and administered sham stimulation (to either the left or right VLPFC, depending on which stimulation group the participant was assigned to) throughout Session 1. Based on each participant’s group assignment, the unblinded experimenter either switched the HD-tDCS device mode to anodal stimulation or pretended to manipulate it but actually left it set to sham stimulation before the participant began Session 2. Although all participants were told that settings were being adjusted, neither the participant nor the blind experimenter who interacted with the participant ever explicitly knew the stimulation mode in either of the two sessions.

Participants were instructed to learn each 30-word list with the expectation that their memory would be tested with free recall immediately after the list was completed, just as it had been on the practice lists. They were informed that a few of the lists might not be followed by immediate free recall tests, but that they would not know whether testing would be required or omitted until after the conclusion of each list. Thus, they were encouraged to be equivalently engaged on the learning of every list. To ensure that the two sessions shared the same behavioral task structure, the first and third list of each session (i.e., L1, L3, L6, and L8) were always tested with free recall, using the same procedures of P1 and P2. For the other lists (L2, L4, L5, L7, L9, and L10), participants were not given any immediate test and could proceed immediately to the subsequent list. Participants were not made aware of this structure and operated with the assumption that post-list recall testing was randomly assigned.

On Day 2, participants returned (18-30 hours after their Day 1 session) for a recognition test that consisted of all the words from the 6 lists that were not tested on Day 1 (180 words in total, with 90 high-value words and 90 low-value words). These 180 words (90 high-value and 90 low-value) were interspersed with 90 lure words that participants had not studied on Day 1. On each trial a single word was displayed on the screen (without any point value), and participants made an old/new judgment using a 6-point confidence scale, with 6 representing “sure old” and 1 representing “sure new.”

To avoid any item or order-specific effects, we employed a constant protocol for the assignment of words to lists and conditions. All participants learned the same subset of words for the practice lists (P1 and P2; 60 words) and the tested-with-recall lists (L1, L3, L6, and L8; 120 words). For these lists, the assignment of high-value and low-value words was counterbalanced across every two participants in each stimulation group. A total of 270 words were selected to be tested with recognition. The assignment was counterbalanced across every six participants in each stimulation group, such that two-thirds (180) of the words were selected for learning (with half assigned high-values and half assigned low-values) on Day 1 (in L2, L4, L5, L7, L8, and L10) and one-third (90) of the words were reserved for use as recognition lures on Day 2. In this way, words that appeared only as lures from some participants were studied by other participants. The presentation orders of these subsets of words were fully randomized during learning on the first day and during recognition testing on Day 2.

### 2.5 Data Exclusion

We excluded nine participants’ data (three in the left-anode group, four in the right-anode group, and two in the sham group) for the following reasons: Two participants had incomplete or corrupted data files due to technical issues. Six participants had extremely poor performance in either the Day 1 recall or Day 2 recognition test. Specifically, we excluded the data of those who recalled fewer than five words on average (i.e., below 16.7% accuracy) on the Day 1 lists (L1, L3, L6, L8) that were tested with immediate free recall (three participants), and/or had an overall *d’* value below 0.25 in the Day 2 recognition task for all 180 words (five participants, two of whom also performed below criterion in the recall task). One additional participant was excluded due to unusually high performance for high-value words in the recognition task, which rendered this participant an extreme outlier whose memory performance (*d’* = 3.59 for the 90 high-value words in both sessions) was 4.1 *SD* above the overall mean.

### 2.6 Data Analysis

For those lists tested with immediate recall on Day 1, both encoding and retrieval processes occurred under the influence of HD-tDCS and could not be dissociated, and thus immediate free recall performance on these lists is not a pure index of how stimulation modulated memory encoding (see Supplementary Materials for reporting on Day 1 free recall performance). On the other hand, because any lingering influences of HD-tDCS on brain function would have worn off well before the Day 2 recognition memory test, we focused our analysis on data from this test (i.e., subsequent memory for words from those lists that were untested on Day 1) to evaluate the effects of stimulation specifically on encoding. Each Day 1 session consists of three untested lists, including 45 high-value words and 45 low-value words. The studied words from Session 1 and Session 2 were intermixed during the Day 2 recognition test, but we were interested in whether memory would differ for words encoded during these two sessions as a function of word value. As the standard deviation of the raw ratings of studied words was significantly larger than that of lure words (the average ratio across 63 participants was 1.21 (*SD* = 0.207) and was significantly greater than 1, *t*(62) = 8.03, *p* < 0.001), and to take full advantage of our 6-point recognition confidence scale (which provides more information about memory strength than old/new judgments alone, we conducted a receiver operating characteristic (ROC) curve analysis using the unequal-variance signal-detection (UVSD) model (Brady et al., 2022; Mickes et al., 2007). This UVSD model yields memory sensitivity values (sometimes denoted as *d*_*a*_) that are analogous to standard *d’* values that could be derived from the hit rates and false alarm rates alone, but are considered a superior measure because their derivation takes into account the entire distribution of confidence judgements to old and new items (Brady et al., 2022). For simplicity, we henceforth use the expression *d’* to refer to these UVSD-derived memory sensitivity estimates. We obtained the *d’* values for high-value (HV) and low-value (LV) words in each session. Note that because the novel lure words presented on the Day 2 memory test were not associated with any particular value condition or session, we used the common distribution of responses to these new items when calculating *d’* for HV and LV items.

The memory *encoding selectivity* (i.e., the value-based memory effect) in each session can be expressed as: HV words *d’* – LV words *d’*. For each individual participant, we subtracted the memory encoding selectivity of Session 1 from that of Session 2, resulting in a difference score that we defined as the overall *encoding selectivity change*:

Encoding selectivity change = Session 2 encoding selectivity (measured by **Δ** *d’*) – Session 1 encoding selectivity (measured by **Δ** *d’*)

To better understand the potential mechanisms underlying the memory encoding selectivity change induced by HD-tDCS, we also employed the dual-process signal-detection (DPSD) model (Yonelinas, 1994, 1999) to dissociate recollection- and familiarity-based responding, and we examined whether memory selectivity change measured by separate parameter estimates of recollection and familiarity differed across stimulation groups. The analytical framework can also be expressed by the equations above, except that the *d’* value is replaced by the parameter estimates for recollection and familiarity.

## 3. Results

### 3.1 Value-related selectivity change measured by *d’*

We first conducted a one-way analysis of variance (ANOVA) of the value-related selectivity change measured by *d’* based on the UVSD model across the three stimulation groups. A significant difference was found across groups, *F*(2, 60) = 5.141, *p* = 0.009, *η*^2^ = 0.146. Post-hoc *t*-tests revealed that the selectivity change measured by *d’* in the left-anode group was significantly larger than that in the right-anode group (Fig. 3, red vs. green bars; post-hoc *t*-test with Holm-Bonferroni correction for multiple comparisons: *p* = 0.011 (uncorrected *p* = 0.004), Cohen’s *d* = 0.908). The selectivity change in the left-anode group was also significantly larger than that in the sham group (Fig. 3, red vs. blue bars; post-hoc *t*-test with Holm-Bonferroni correction: *p* = 0.034 (uncorrected *p* = 0.017), Cohen’s *d* = 0.700). The right-anode group and the sham group did not differ in the overall selectivity change, *p* = 0.540 (uncorrected *p* = 0.540), Cohen’s *d* = −0.216. This is evidence that anodal stimulation of left VLPFC resulted in higher overall memory encoding selectivity, while this effect was not observed for the contralateral site in the right VLPFC.

**Figure 3.**
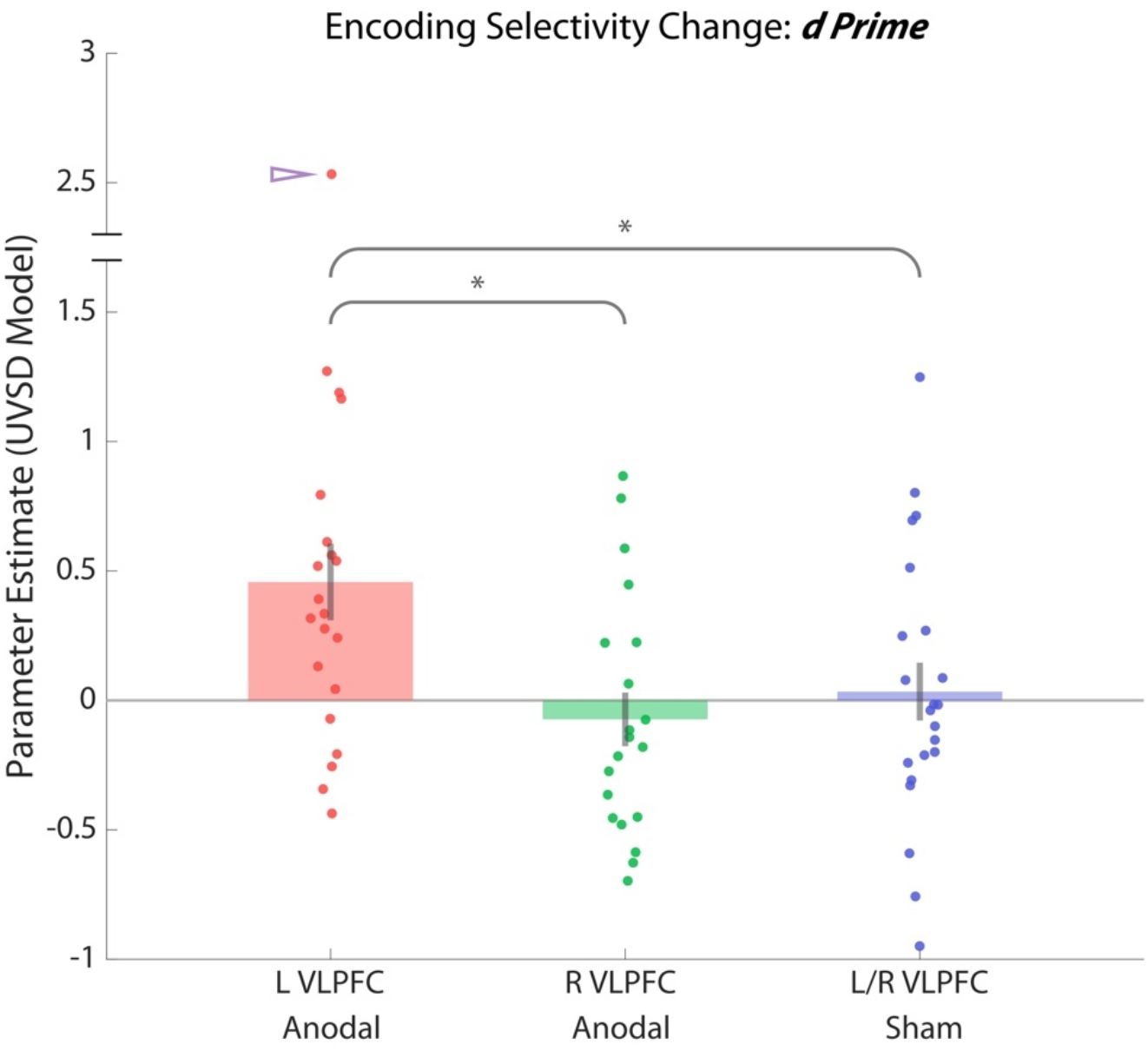
Memory encoding selectivity change from Session 1 to Session 2 measured by *d’* under the UVSD model. A significant difference in memory encoding selectivity change measured by *d’* was found across stimulation groups. Post-hoc analyses revealed that the memory encoding selectivity change in the left-anode group was significantly larger than that in the right-anode group and the sham group. Each dot represents the data from an individual participant. The purple triangular pointer indicates a participant in the left-anode group who exhibited particularly high memory selectivity change (note the discontinuity in the y-axis values). Although this participant was not technically an outlier according to our criteria, see main text for results when data were re-analyzed without this participant. \**p* < 0.05 (Holm-Bonferroni corrected for multiple comparisons)

It is possible that the effects observed above were heavily influenced by the one participant in the left-anode group (marked with the purple triangular pointer in Fig. 3) who exhibited dramatic positive change in memory selectivity for high-value words (1.14 of *d’* change across sessions) and negative change in memory selectivity for low-value words (−1.40 of *d’* change across sessions). Although this participant did not constitute as an outlier under any of the data exclusion criteria, we also examined the results without this data point to ensure that the group-level differences were not driven by the outstanding encoding selectivity change of this single subject. When we performed the analyses above while excluding this participant, the core group effects remained, although their effect size was slightly diminished. The ANOVA again revealed a significant difference in memory selectivity change across groups, *F*(2, 59) = 4.013, *p* = 0.023, *η*^2^ = 0.120. The difference between the left-anode group and right-anode group remained significant (post-hoc *t*-test with Holm-Bonferroni correction for multiple comparisons: *p* = 0.026 (uncorrected *p* = 0.009), Cohen’s *d* = 0.888). The comparison between the left-anode group and sham group only trended towards significance after Holm-Bonferroni correction but still yielded a medium-to-large effect size, *p* = 0.084 (uncorrected *p* = 0.042), Cohen’s *d* = 0.624.

### 3.2 Value-related selectivity change measured by recollection and familiarity

We then sought evidence for process dissociations in the effects of HD-tDCS on value-related memory selectivity. We did this by computing the parameter estimates for recollection and familiarity derived by applying the DPSD model separately to recognition confidence ratings for high-value and low-value studied words that had appeared in Session 1 or Session 2, using ratings for unstudied lures as the common basis for the false alarm rate component of the curves. All curve fitting and parameter estimation was accomplished using the ROC Toolbox (Koen et al., 2017). The resulting recollection and familiarity estimates were then subjected to statistical analysis using the same framework applied above.

We found a significant value-related selectivity change on recollection-based memory across the three stimulation groups using a one-way ANOVA, *F*(2, 60) = 3.652, *p* = 0.032, *η*^2^ = 0.109. Post-hoc *t*-tests revealed that the selectivity change measured by recollection in the left-anode group was significantly larger than that in the right-anode group (Fig. 4A, red vs. green bars; post-hoc *t*-test with Holm-Bonferroni correction for multiple comparisons: *p* = 0.044 (uncorrected *p* = 0.015), Cohen’s *d* = 0.703). The selectivity change in the left-anode group was also larger than that in the sham group (Fig. 4A, red vs. blue bars), though this comparison only trended towards significance after Holm-Bonferroni correction for multiple comparisons, *p* = 0.076 (uncorrected *p* = 0.038), Cohen’s *d* = 0.672. We did not find any value-related selectivity change on familiarity-based memory across the three stimulation groups (Fig. 4B), *F*(2, 60) = 0.250, *p* = 0.779, *η*^2^ = 0.008. Our results thus suggest that anodal stimulation of left VLPFC primarily boosted the impact of item value on the encoding of memories that lead to subsequent recollection.

**Figure 4.**
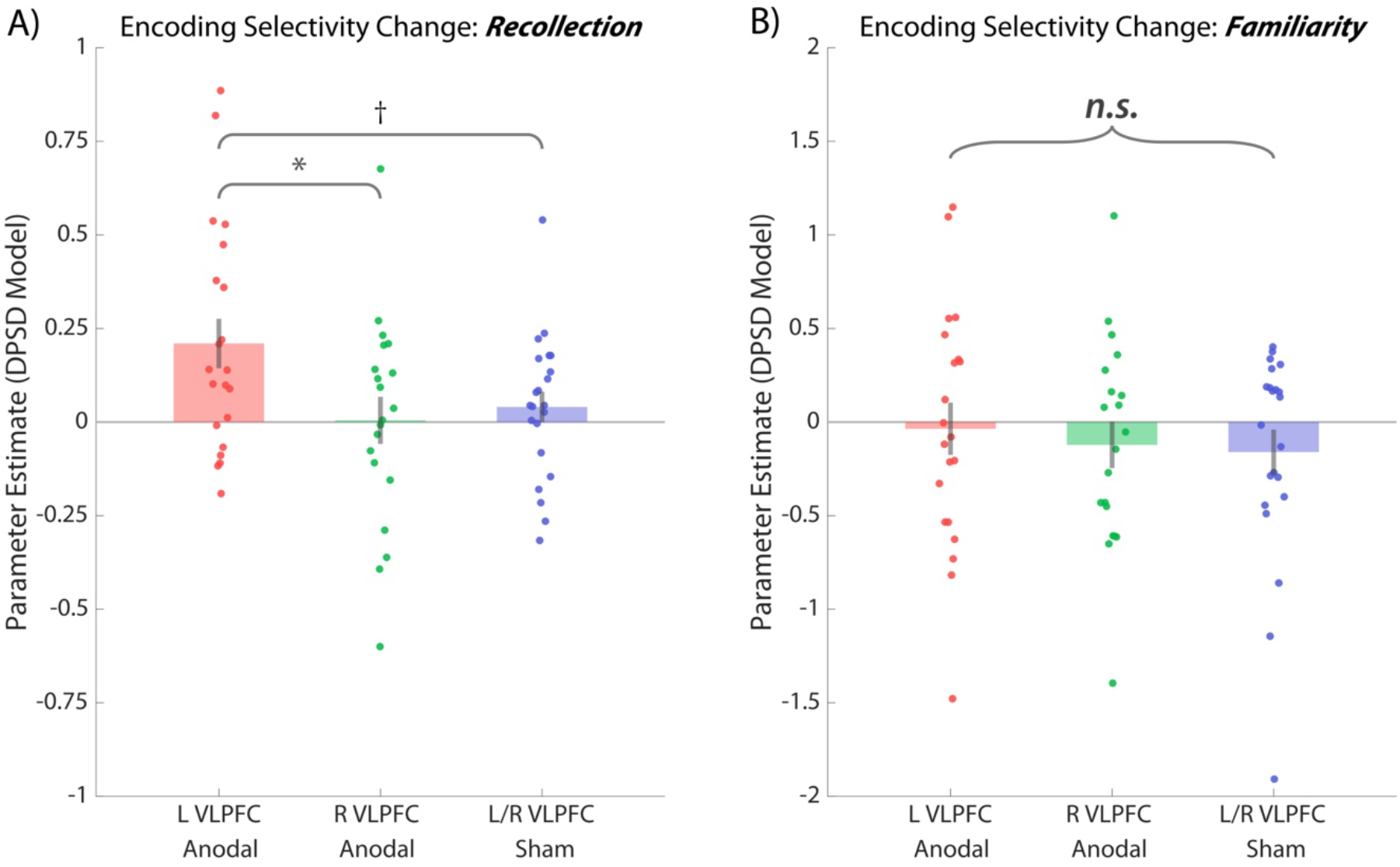
Memory encoding selectivity change from Session 1 to Session 2 measured by recollection and familiarity under the DPSD model. **A) Encoding selectivity change measured by recollection**. A significant difference in memory encoding selectivity change measured by recollection was found across stimulation groups. Post-hoc analyses revealed that the encoding selectivity change in the left-anode group was significantly larger than that in the right-anode group, but the difference between the left-anode group and sham group only trended towards significance. **B) Encoding selectivity change measured by familiarity**. No significant difference in encoding selectivity change measured by familiarity was found across stimulation groups. Each dot represents the data from an individual participant. \**p* < 0.05, †*p* < 0.10 (Holm-Bonferroni corrected for multiple comparisons)

## 4. Discussion

Previous neuroimaging studies have shown that fronto-temporal semantic processing regions (especially the left VLPFC) are heavily activated during the encoding of high-value items relative to low-value items, and the degree of their engagement correlates with individual differences in memory selectivity (Cohen et al., 2014, 2016). In the present study, we established causal evidence for the role of left VLPFC in value-directed remembering. Specifically, we found that targeted anodal stimulation of the left VLPFC accentuates the impact of item value on subsequent recognition memory, assessed one day after stimulation. Using a double-blind stimulation protocol and two control groups, we showed that the increase in value-directed memory encoding selectivity caused by anodal stimulation of left VLPFC is not observed when stimulation is applied to the homologous region of right VLPFC, nor when sham stimulation was applied. This provides compelling evidence for the left-lateralization of VLPFC contributions to selective encoding, at least as it pertains to the memorization of word stimuli. While memory encoding almost certainly engages both hemispheres and depends on communication between them, the differences between left and right stimulation demonstrate hemispheric processing biases (Zaidel, 1983) in verbal memory encoding.

What are the mechanisms by which increased cortical excitability of left VLPFC leads to more selective and effective encoding? We speculate that left VLPFC may facilitate the encoding of high-value words by linking them to broader semantic information, such as a narrative context (i.e., using the words to make up a story), creating a meaningful mental image of the item, or a self-referential context (i.e., thinking about how the words relate to oneself or one’s past experiences). Because we did not observe an overall enhancement of memory encoding with left VLPFC stimulation, it appears that stimulation made left VLPFC more effective at fostering elaborative semantic encoding for those items selected as important. This interpretation fits with the idea that anodal tDCS does not cause neurons to fire but instead brings them closer to their firing threshold, such that they would be more likely to exceed their threshold and fire when engaged in a cognitive process. It is important to note that the relevant semantic information might not be primarily generated by the region of the left VLPFC that we targeted. The mid-VLPFC region that received the greatest current intensity in our stimulation protocol (i.e. Brodmann Area 45) is more likely to play a role in selecting which of the many pieces of semantic knowledge evoked by each word one should attend to and further elaborate on (Badre & Wagner, 2007; Thompson-Schill et al., 1997). The semantic retrieval process itself may be mediated by other functionally connected regions such as the pre-supplementary motor area (pre-SMA; Crosson et al., 2001; Hart et al., 2013; Lou et al., 2017), the lateral temporal cortex (Binder et al., 2009; Bonner & Price, 2013), and more anterior regions of left VLPFC, such as Brodmann Area 47 (Badre & Wagner, 2007).

Another possible interpretation of the beneficial effects of anodal left VLPFC stimulation is that it facilitated cognitive control mechanisms such that participants were better able to apply differential encoding based on item value. The stimulation may have enhanced metacognitive strategies about which items to effectively encode and which items to avoid encoding, as VLPFC is indeed involved in the formulation and selection of high-level strategies that guide attention (Hampshire & Owen, 2005; Koechlin et al., 2003; Owen & Hampshire, 2009). In the present study, we found that the stimulation of the left VLPFC resulted in not only enhanced encoding of high-value words but also reduced encoding of low-value words (see Supplemental Materials, section 2). Although the stimulation-induced reduction in subsequent memory for low-value words did not achieve statistical significance and thus should be interpreted with caution, it is generally consistent with an effect of stimulation on metacognitive control. Thus, we cannot fully disentangle the semantic component from the strategic component with our current findings, and future studies may be necessary to address the distinctive role of each component in value-directed remembering. However, the clearly left lateralized effect of stimulation we observed is more consistent with an effect on semantic processing than cognitive control, as cognitive control typically involves bilateral involvement of prefrontal regions (Miller & Cohen, 2001; Ryman et al., 2019) and thus should have been enhanced by right VLPFC stimulation as well.

It is also possible that anodal left VLPFC stimulation enhanced dopaminergic reward-based learning of high-value items. Prior fMRI experiments show that reward-sensitive regions, such as the ventral striatum and midbrain regions, also exhibit sensitivity to value in the VDR paradigm (Cohen et al., 2014). Furthermore, the dopaminergic reward system might interact with the prefrontal areas to guide motivational and controlled behaviors (Arias-Carrión et al., 2010; Brenhouse et al., 2008; Telzer, 2016; Telzer et al., 2015). Thus, reward-based learning may also be engaged in prioritizing the memorization of high-value relative to low-value words in the VDR paradigm. The fact that significant effects of stimulation were seen in recollection and after a one-day delay is consistent with the facilitation of reward modulation of hippocampal-dependent episodic encoding (Adcock et al., 2006; Shohamy & Adcock, 2010). However, the left-lateralized effect of tDCS seen here is less consistent with this interpretation, as the dopaminergic reward system is bilateral and the anatomical route by which anodal VLPFC stimulation is activating this system is less clear.

Stimulation of left VLPFC significantly increased memory encoding selectivity measured by recollection but did not reliably enhance selectivity on a measure of familiarity. Previous work has consistently shown enhanced recollection of high-value items in the context of value-directed remembering, but enhancement of familiarity for high-value items has been more equivocal (Cohen et al., 2017; Elliott et al., 2020; Hennessee et al., 2017). The present work suggests that VLPFC stimulation induced encoding of high-value words in a manner that led to distinctive memory traces that would have a greater likelihood of recollection later. Since our measures of recollection and familiarity were estimated based on parameters derived from ROC curves fitted on confidence rating data rather than asking participants to make Remember/Know judgments, we cannot be sure that participants had the subjective experience of recollection per se, but stronger recollection estimates suggest that participants were highly accurate when indicating the highest level of recognition confidence. Cohen et al. (2017) found effects of value on both recollection and familiarity when participants were given the opportunity to use metacognitive control to modify their memory encoding strategies across the study-test cycles through recall tests with feedback, similar to the present study, and suggested that increased strategic encoding of high-value items led to greater general strength of these items across both recollection and familiarity. While the present study did not replicate that result, one explanation is that recognition in the present study was tested after a one-day delay, which may have reduced effects on familiarity. Still, we were able to identify enhanced recollection-based selectivity induced by anodal stimulation of left VPLFC, perhaps because deep semantic encoding processes such as self-referential processing enrich the context associated with the words, allowing for conscious recollection of more episodic details during retrieval.

In the present study, we incorporated several controls that helped with isolating the effect of HD-tDCS on memory encoding selectivity. We incorporated a within-subject comparison in our design, with every subject undergoing sham stimulation in the first session on Day 1 where their performance served as the baseline. This reduces the impact of individual variability in verbal encoding selectivity to a great extent. The individual differences in encoding selectivity may lessen the effect sizes of HD-tDCS if we implemented a solely between-subject design, given the huge across-participant variation in objectively and subjectively measured encoding strategies and the resulting selectivity demonstrated in the VDR paradigm (Cohen et al., 2017). With the within-subject across-session subtraction of baseline performance, we were able to emphasize the effect of HD-tDCS *per se*. Furthermore, we also incorporated controls to isolate memory encoding from the retrieval process. To fully dissociate the two processes, we included several word lists that were not immediately tested on Day 1 but were later tested on Day 2, when HD-tDCS effects no longer remained.

Our findings add to the growing body of tDCS literature that has examined the role of the left prefrontal cortex (PFC) in verbal memory encoding (see Galli et al., 2019, for a review). While prior work has demonstrated that anodal stimulation of the left dorsolateral PFC (DLPFC) enhances verbal working memory (Naka et al., 2018), long-term episodic memory (Javadi & Cheng, 2013; Javadi & Walsh, 2012), and memory monitoring (i.e. metamemory) processes (Chua & Ahmed, 2016), our results are the first neuromodulation evidence that the left VLPFC contributes to the strategic selection and prioritized encoding of valuable information, rather than to memorization of items more generally. Thus when our findings are considered alongside this prior tDCS work, the pattern of results fits nicely with conceptualizations of the differential roles of DLPFC and VLPFC in memory encoding (Blumenfeld & Ranganath, 2006, 2007): while the DLPFC putatively regulates the formation of long-term episodic memory through storing and organizing item-level information in working memory, the VLPFC underlies the development and execution of encoding strategies, selection of goal-relevant items, and optimization of memory encoding efficiency under a limited capacity.

There are several limitations of the present study. First, our findings may be statistically underpowered to observe some effects of stimulation. We ran a posthoc power analysis in G-power (Faul et al., 2007) and found that we achieved 81% and 61% of power for our two major comparisons (*d’* measures: left-anode vs. right-anode and left-anode vs. sham), with the latter comparison slightly below the satisfactory power level. Second, as we only used verbal stimuli (i.e., words) in our experiment, it is unknown if our findings would generalize to non-verbal stimuli r. In fact, Cohen and colleagues (2019) examined value-directed encoding of abstract visual images and found that activity in bilateral VLPFC was associated with successful encoding, especially of high-value images; this activity was more widespread in the left hemisphere, but was also reliably apparent in the right hemisphere. Thus, it is possible that stimulation of the right VLPFC also modulates memory selectivity for non-verbal stimuli. Future work will be necessary to better understand the relative contributions of the left and right VLPFC in the encoding of verbal and non-verbal stimuli. Finally, our results suggest potential interventional applications that may help with improving memory selectivity and efficiency in the short and long run. Many studies have revealed the potential of using neuromodulation techniques to enhance cognitive functioning in patients and healthy individuals, including those cognitive abilities specific to the memory and language domains (see Brunoni et al., 2012, for a review). Crucially, anodal stimulation of the left PFC has been demonstrated to benefit verbal memory in both young adults and older adults who experience an age-related cognitive decline (Manenti, 2016; Manenti et al., 2013; Sandrini et al., 2014, 2016). Our findings demonstrate that while focal neurostimulation of the left VLPFC does not enhance overall memory accuracy, it does seem to facilitate the application of efficacious deep semantic encoding strategies, which maximizes the utility of memory given a constrained overall capacity and the need to prioritize the most important information. Importantly, though our sample consisted of only young adults, we expect that our HD-tDCS intervention could be particularly effective in older adults, as aged individuals tend to engage semantic encoding strategies to an equivalent or larger extent because of their decline in overall memory capacity (Cohen et al., 2016). It may also be fruitful to examine potential applications to populations with impaired value-directed remembering, such as attention-deficit/hyperactivity disorder (Castel et al., 2011) and Alzheimer’s disease (Castel et al., 2009; Wong et al., 2019). While our current study was not designed to assess the possibility of boosting left VLPFC function in an enduring manner that could help people better encode valuable information encountered in their daily lives, future research with multi-day stimulation protocols that foster longer-lasting cortical plasticity might show potential for promoting more effective selective and thus more efficient learning.

## Acknowledgements

We sincerely appreciate Gabriel Damon Lavezzi, Xin Luo, Ashley Ngo, William Stonehouse, Ayona Sengupta, and Ruiwen Zhou (names ordered alphabetically), for their dedicated effort in conducting the experiment and collecting the data. We also thank Guangzheng Xu for his assistance on making some of the figures. This work was supported by the Tsinghua University Initiative Scientific Research Program (20197020005 to L.T.H.) and the US National Science Foundation Grant (NSF BCS-2048692; PI: Barbara Knowlton).

As colleagues of Eran Zaidel at UCLA, we appreciated Eran’s many thoughtful comments and engaging discussions regarding laterality and cognitive functions over the years, and he will be fondly remembered for his generosity, wisdom, and creativity.

