## Supplemental Materials for "Establishing a causal role for left ventrolateral prefrontal cortex in value-directed memory encoding with high-definition transcranial direct current stimulation"

1. **Performance on Day 1 in immediate free recall tests**

On Day 1, each session consists of two lists that were tested with immediate free recall, after which participants were given instant feedback on their performance. Here, we present the results on participants’ free recall performance in each stimulation group. The numbers of words recalled are presented in Fig. S1A, with high-value (HV) words and low-value (LV) words plotted separately. High-value words were significantly more frequently recalled in all the lists in all the stimulation groups (all *P*s < 0.01). Then, we subtracted the number of LV words recalled from that of HV words and plotted the difference in Fig. S1B. This figure represents the selectivity in recall across all the lists. Similar to our approach of computing the memory encoding selectivity change across sessions, we examined whether the selectivity change in recall from Session 1 (L1 and L3) to Session 2 (L6 and L8), measured by the following equation, differed across the three stimulation groups:

Selectivity change = Session 2 selectivity (measured by # HV words - # LV words) - Session 1 selectivity (measured by # HV words - # LV words)

The one-way analysis of variance (ANOVA) revealed no significant difference in the selectivity change in recall across the three stimulation groups, *F*(2, 60) = 0.733, *p* = 0.485, *η*^2^ = 0.024.

Next, we analyzed a different measure of how value impacts recall, known as the *selectivity index (SI)*. This is a measure of recall selectivity that takes into account the participant’s actual obtained score on each list relative to the maximum score that could have been obtained given the number of words recalled. This was calculated with the following equation for each list (Castel et al., 2002):

$$SI= \frac{actual score - chance score}{ideal score - chance score}$$

Given our paradigm, the chance score is the average value of all the words in each list (6.5) multiplied by the number of words recalled. The ideal score depends on the number of words recalled; for example, if one recalled 17 words out of a 30-word list, then the maximum possible score (i.e., ideal score) would be 5 × 12 + 5 × 11 + 5 × 10 + 2 × 3 = 171. In this case, if the participant obtained 130 points in total, then the SI would be (130 – 6.5 × 17)/(171 - 6.5 × 17) = 0.322. The SI of each session was calculated by averaging the selectivity index of the two lists in that session.

Because we were interested in how SI changed across the two sessions, we subtracted the mean SI from Session 1 from the mean SI from Session 2. The mean SI change scores (± SD) for each group were as follows: left-anode, 0.119 (± 0.263); right-anode, 0.097 (± 0.274); sham, 0.043 (± 0.270). A one-way ANOVA showed that the selectivity change was not significantly different across the three stimulation groups, *F*(2, 60) = 0.461, *p* = 0.633, *η*^2^ = 0.015.

Though our results showed that anodal stimulation of the left VLPFC did not significantly enhance free recall selectivity, we believe that immediate free recall is a less sensitive measure compared with recognition in terms of reflecting the memory strength of all the encoded words in our experimental paradigm. First, we incorporated only two tested-with recall lists but three tested-with recognition lists, which renders the free recall data underpowered. Second, successful recall of a given item requires that item to surpass a certain threshold of memory strength. Take high-value (HV) words as an example – even if the memory strength of all HV words has increased by some amount due to anodal tDCS of the left VLPFC, only a small portion of the items will exceed the recall threshold. Perhaps one extra HV word would be recalled, despite the fact that all of them may have received a boost in strength. This boost in strength is thus better detected on a recognition test that queries memory strength for all of the words. This is probably why we did not observe a significant across-group difference in the free recall data. Furthermore, note that all the recall procedures underwent HD-tDCS, so the effects of HD-tDCS on memory encoding cannot be isolated from those on memory retrieval.

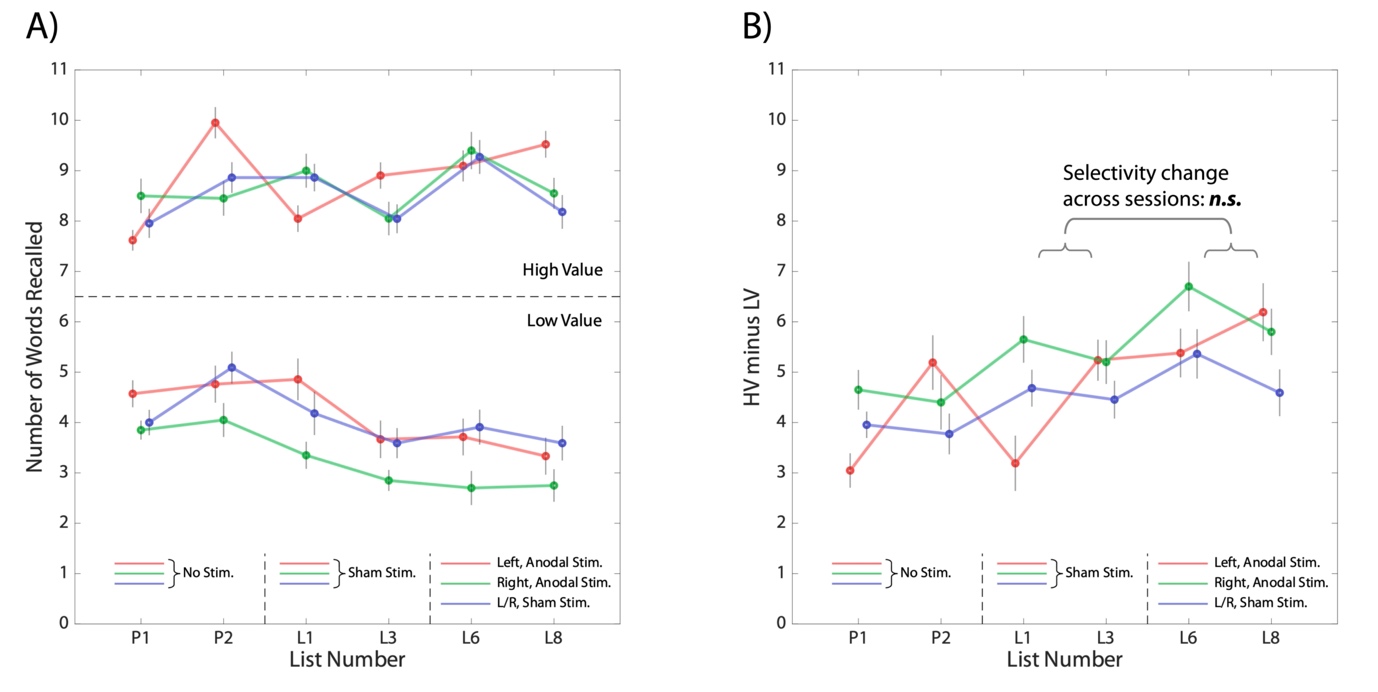

**Figure S1. Recall performance (in lists tested on Day 1) in each stimulation group.**

**A) Recall performance in all the six lists tested with recall.** Numbers of HV words and LV words successfully recalled in each list are plotted separately. HV words were recalled more frequently than low-value words in all the lists in all the stimulation groups. **B) Memory selectivity in all of the six lists tested with recall.** Memory selectivity in each list was computed by the difference in the number of HV and LV words recalled. The selectivity change from Session 1 (L1 and L3) to Session 2 (L6 and L8) was not significantly different across the three stimulation groups.

2. **The effects of HD-tDCS on the overall memory performance**

We examined whether HD-tDCS improved the overall recognition memory performance. We calculated the *d’* under the UVSD model for all the words (HV and LV combined) in each session and examined the overall memory performance change from Session 1 to Session 2. The results are shown in Table S1. None of the three groups exhibited positive change in the overall memory performance (likely due to decreased encoding of LV words), and there was no significant difference between the three stimulation groups, *F*(2, 60) = 0.912, *p* = 0.407, *η*^2^ = 0.030. This is evidence that the stimulation of the left VLPFC did not improve the overall memory performance while enhancing memory encoding selectivity.

**Table S1 Overall memory performance in the two sessions**

| Stimulation Group | *d’* for all the words (HV and LV combined) | |  | *d’* Change |
| --- | --- | --- | --- | --- |
|  | Session 1 | Session 2 |  |  |
| Left-anode (*N* = 21) | 1.04 (± 0.37) | 0.90 (± 0.38) |  | -0.13 (± 0.19) |
| Right-anode (*N* = 20) | 0.99 (± 0.51) | 0.94 (± 0.42) |  | -0.05 (± 0.27) |
| Sham (*N* = 22) | 1.12 (± 0.56) | 0.95 (± 0.48) |  | -0.16 (± 0.35) |

3. **Dissecting the effects of HD-tDCS on memory for high- and low-value words**

In the main text, we reported the memory selectivity change across the two sessions, which is computed as (Session 2 HV memory sensitivity – Session 2 LV memory sensitivity) – (Session 1 HV memory sensitivity – Session 1 LV memory sensitivity). Here, we also report in Table S1 each stimulation group’s original mean and standard deviation values of *d’* (UVSD model) for HV and LV words in the two sessions.

**Table S2** **Memory (d') for HV and LV Words in the Two Sessions (M ± SD)**

| Stimulation Group | High-value words | |  | Low-value words | |
| --- | --- | --- | --- | --- | --- |
|  | Session 1 | Session 2 |  | Session 1 | Session 2 |
| Left-anode (*N* = 21) | 1.30 (± 0.48) | 1.45 (± 0.57) |  | 0.82 (± 0.48) | 0.52 (± 0.49) |
| Right-anode (*N* = 20) | 1.41 (± 0.66) | 1.33 (± 0.64) |  | 0.63 (± 0.51) | 0.62 (± 0.42) |
| Sham (*N* = 22) | 1.47 (± 0.62) | 1.32 (± 0.63) |  | 0.83 (± 0.66) | 0.65 (± 0.46) |

Next, to further explore the mechanisms underlying the memory selectivity change induced by HD-tDCS, we examined the change in encoding of high-value and low-value words separately, as the overall selectivity change measures considered above could be ascribed to enhanced encoding of high-value words, reduced encoding of low-value words, or a combination of both effects. We computed the memory encoding change for HV and LV words separately with the following equation:

HV/LV words encoding selectivity change = Session 2 HV/LV words *d’* – Session 1 HV/LV words *d’*

We first conducted a two-way mixed ANOVA (stimulation type × value) to examine the effect of stimulation and value on memory change measured by *d’* (Fig. S2A; this is the same analysis with the one-way ANOVA for Section 3.1 in the main text). A significant interaction was revealed, *F*(2, 60) = 5.141, *p* = 0.009, *η*^2^ = 0.146. We then analyzed the simple effect of stimulation for high-value and low-value words separately. For high-value words, the main effect of stimulation only trended towards significance after Holm-Bonferroni correction, *F*(2, 60) = 2.847, *p* = 0.066 (uncorrected *p* = 0.066), *η*^2^ = 0.087. The enhancement effect for high-value words in the left-anode group was numerically larger than that in both the right-anode group and the sham group, but this was not statistically significant under the Holm-Bonferroni correction (left-anode vs. right-anode: Holm-Bonferroni corrected *p* = 0.176 (uncorrected *p* = 0.088), Cohen’s *d* = 0.565; left-anode vs. sham: Holm-Bonferroni corrected *p* = 0.077 (uncorrected *p* = 0.026), Cohen’s *d* = 0.695). For low-value words, a significant main effect of stimulation was found, *F*(2, 60) = 3.947, *p* = 0.049 (uncorrected *p* = 0.025), *η*^2^ = 0.116. Memory for low-value words in the left-anode group was significantly reduced compared with the right-anode group (Holm-Bonferroni corrected *p* = 0.021 (uncorrected *p* = 0.007), Cohen’s *d* = -0.990), yet was not statistically different from the sham group (Holm-Bonferroni corrected *p* = 0.235 (uncorrected *p* = 0.235), Cohen’s *d* = -0.331).

We then applied the same two-way mixed ANOVA pipeline to analyzing the recollection and familiarity data. We found a significant interaction between stimulation type and value for recollection (Fig. S2B) but not familiarity (Fig. S2C), which was exactly the same as the results when the difference between high-value and low-value words was calculated, as reported in the main text. Next, we analyzed the simple effect of stimulation on recollection for high-value and low-value words. For high-value words, the main effect of stimulation on recollection only trended toward significance under Holm-Bonferroni correction, *F*(2, 60) = 3.789, *p* = 0.056 (uncorrected *p* = 0.028), *η*^2^ = 0.112. The enhancement in recollection for high-value words in the left-anode group was significantly larger than that in the right-anode group (Holm-Bonferroni corrected *p* = 0.045 (uncorrected *p* = 0.015), Cohen’s *d* = 0.698). It was also larger than that in the sham group, although this only trended towards significance after Holm-Bonferroni correction (Holm-Bonferroni corrected *p* = 0.058 (uncorrected *p* = 0.029), Cohen’s *d* = 0.717). For low-value words, the main effect of stimulation was not significant, *F*(2, 60) = 1.200, *p* = 0.308, *η*^2^ = 0.038.

While our core analyses in the main text focused on high-minus-low-value difference scores, when we separately examined memory for high- and low-value words, we saw modest evidence for both a stimulation-induced elevation of memory for high-value words and a diminution of memory for low-value words. That said, statistical analysis of these isolated effects was complicated by reduced power, and some of the contrasts did not achieve significance when corrected for multiple comparisons. This makes it challenging for us to confidently conclude that anodal stimulation of left VLPFC causes both a memory enhancing effect for high-value items and a memory suppressing effect for low-value items (e.g., by actively inhibiting the processing of these items or withdrawing attention from them to instead prioritize rehearsal of other recently presented high-value items). Rather, our data lead us to a more neutral conclusion that stimulation of this region boosted memory selectivity as a function of item value. Future studies with greater power may be necessary to determine the relative contribution of enhancement and suppression mechanisms. But critically, none of these value-related memory effects were observed in the control groups that received right VLPFC or sham stimulation.

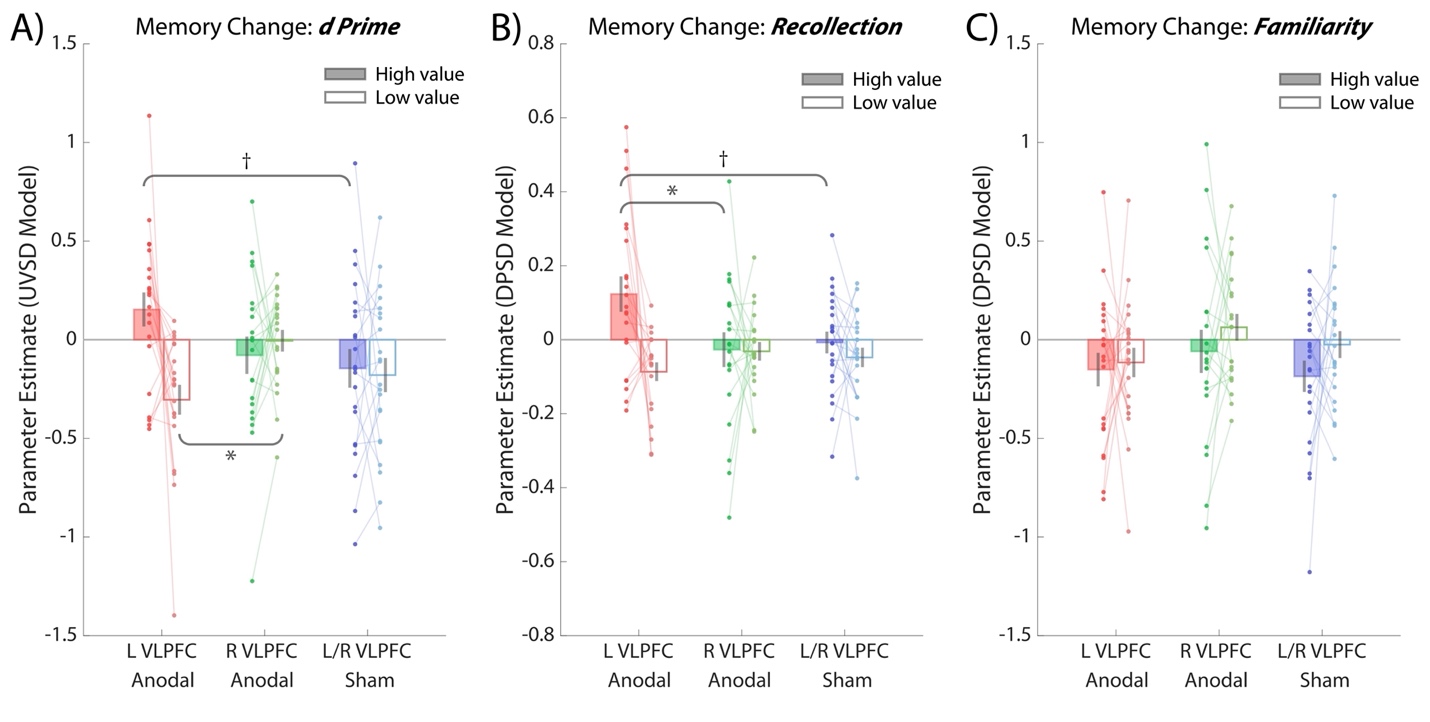

**Figure S2. Memory change from Session 1 to Session 2, with high-value and low-value words analyzed separately.**

**A) Change in recognition memory (as measured by *d’*) for HV and LV words.** There was a significant interaction of stimulus value and group membership on memory change. For high-value words (filled bars), the effect of group only trended towards significance. For low-value words (unfilled bars), there was a significant main effect of group, yet post hoc tests revealed that only the difference between the left-anode group and right-anode group achieved significance. **B) Change in recollection for HV and LV words.** There was a significant interaction of stimulus value and group membership on recollection change. For high-value words, the effect of group only trended towards significance. The difference between the left-anode and right-anode group was statistically significant, yet the difference between the left-anode and sham group only trended towards significance. For low-value words, no significant difference across groups was revealed. **C) Change in familiarity for HV and LV words.** There was no interaction of stimulus value and group membership on familiarity change. For both high-value and low-value words, no main effect of stimulation on familiarity was found. Each dot represents the data from an individual participant, with connected data points illustrating value-related changes within-subjects.

^*^*p* < 0.05, ^†^*p* < 0.10 (Holm-Bonferroni corrected for multiple comparisons); non-significant results are not indicated in the figure.
